# Environment affects specialisation of plants and pollinators

**DOI:** 10.1101/866772

**Authors:** E. Fernando Cagua, Audrey Lustig, Jason M. Tylianakis, Daniel B. Stouffer

**Affiliations:** Centre for Integrative Ecology, School of Biological Sciences, University of Canterbury, Private Bag 4800, Christchurch 8041, New Zealand; Geospatial Research Institute, University of Canterbury, Private Bag 4800, Christchurch 8041, New Zealand

**Keywords:** eltonian niche, environmental effects, generalisation & specialisation, species degree, species interactions, throphic niche

## Abstract

What determines whether or not a species is a generalist or a specialist? Evidence that the environment can influence species interactions is rapidly accumulating. However, a systematic link between environment and the number of partners a species interacts with has been elusive so far. Presumably, because environmental gradients appear to have contrasting effects on species depending on the environmental variable. Here, we test for a relationship between the stresses imposed by the environment, instead of environmental gradients directly, and species specialisation using a global dataset of plant-pollinator interactions. We found that the environment can play a significant effect on specialisation, even when accounting for community composition, likely by interacting with species’ traits and evolutionary history. Species that have a large number of interactions are more likely to focus on a smaller number of, presumably higher-quality, interactions under stressful environmental conditions. Contrastingly, the specialists present in multiple locations are more likely to broaden their niche, presumably engaging in opportunistic interactions to cope with increased environmental stress. Indeed, many apparent specialists effectively behave as facultative generalists. Overall, many of the species we analysed are not inherently generalist or specialist. Instead, species’ level of specialisation should be considered on a relative scale depending on where they are found and the environmental conditions at that location.

## Introduction

Species interactions are known to vary widely across space and time. There are multiple examples of species that interact with a large number of partners in a particular community or season, but with fewer in another. Some of this variation can be attributed to environmental drivers. However, how exactly the environment, specifically the stress it imposes on species, affects whether two species interact or not, and ultimately the species’ specialisation. Understanding how the environment drives the number of partners is crucial because it underpins the species’ role in its community and shapes the structure of the network of interactions. This structure, in turn, determines ecosystem function and stability.

Species interactions are determined in part by niche processes (the matching of traits) and partly by neutral processes (more abundant species are more likely to encounter each other and, thus, interact). The environment can influence both of these processes. It is, therefore, not surprising that, despite limitations on the spatial extent or the number of environmental gradients considered, multiple studies have been able to show how changes to interactions can be related to environmental change (Tylianakis and Morris 2017). For instance, some studies suggest that the strength of some trophic interactions, like predation (McKinnon et al. 2010; Vucic-Pestic et al. 2011) and herbivory (Baskett and Schemske 2018), can increase with temperature but might decrease with precipitation (Pires et al. 2016). Some other studies, however, have shown either no effect (on average) or non-linear effects of temperature or precipitation on plant-pollinator interactions (Devoto, Medan, and Montaldo 2005; Gravel et al. 2018). Overall, while it looks clear that pairwise interactions respond to environmental drivers, there is high variability in the response (Tylianakis et al. 2008).

One possible explanation for the seemingly contradictory evidence is that different bioclimatic factors (like temperature or precipitation) can have contrasting effects on species and their partners. Here we attempt to simplify this situation by reducing multiple factors into a single measure of environmental stress. Previous research suggests that environmental stress may affect the number of partners in different ways depending on its role in the community (for example its trophic guild) or even the species itself. Specifically, we propose two alternative hypotheses of how environmental stress may affect specialisation (Tylianakis and Morris 2017). On the one hand, it is possible that when species are under environmental stress, they might be “pressured” to focus on partners with which they are best adapted to interact. For instance, Hoiss et al. (2012) found increased phylogenetic clustering between plants and pollinators at higher altitudes; while Peralta et al. (2015) found that parasitoids in plantation forest, where environmental stress was higher than in native forests, were constrained to interact with hosts, they were best adapted to attack. Similarly, Lavandero and Tylianakis (2013) showed that environmental stress due to higher temperature reduced the trophic niche breadth of parasitoids suggesting higher specialisation.

On the other hand, it is also possible that when species are under environmental stress, they are forced to be more flexible in their interactions. Higher environmental stress is likely to be reflected in greater energetic or reproductive costs. Therefore they might not be able to sustain encounter rates with their preferred partners at sufficient levels. In line with this hypothesis, Hoiss, Krauss, and Steffan-Dewenter (2015) found that the specialisation of plant-pollinator networks decreased both with elevation and after extreme drought events. Likewise, Pellissier et al. (2010) found a positive relationship between niche breadth and environmental stress: disk- or bowl-shaped blossoms (which allow a large number of potential pollinator species to access pollen and nectar rewards) dominated at high altitude flower communities.

Here, we investigate whether and how environmental stress can systematically affect specialisation. Our main aim is to test the two hypotheses mentioned above that relate environmental stress and species’ number of partners and investigate whether this changes across species or between trophic guilds. We propose that specialist species can become “facultative” generalists to reduce their vulnerability to the absence of preferred partners (for example, when variations in climate decouple phenologies; Benadi et al. 2014). In other words, we expect that, as environmental stress increases, specialists should be more likely to engage with more partners. Species with a large number of partners, on the other hand, should have a larger pool of available partners and might, therefore, be more likely to specialise under environmental stress and focus on the most beneficial partners. Importantly, when testing these hypotheses, we control for the potential effects of the environment in community composition (which has been previously shown to be a determinant factor; Gravel et al. 2018). We test these hypotheses using data on plant-pollinator interactions. These interactions provide a particularly interesting system to test these hypotheses because, due to the multiple trade-offs involved in the pollination service, there are multiple intuitive ways in which we could imagine species respond to environmental stress given the available partners. We estimate the stress species might experience in their community by calculating the bioclimatic suitability of their communities given the species’ patterns of global occurrence.

## Methods

We retrieved plant-pollinator networks from the Web of Life database (Fortuna, Ortega, and Bascompte 2014). This database contains datasets originating from 57 studies published in the primary literature between 1923 and 2016. Calculating the environmental stress of species in their community and their potential partners required us to reduce both the taxonomic and distributional/locational uncertainty. A critical step towards reducing this uncertainty is to ensure that the names used to identify species are valid and unambiguous, which in turn allow us to obtain further information from biological databases and accurately match species across studies. Therefore, our first step was to ensure consistent spelling and standardisation of species names synonyms (see Supplementary Methods). The cleaning process resulted on a total of 2,555 plants and 8,406 pollinator species distributed across 73 locations arround the globe (Figure 1 and S1).

**Figure 1:**
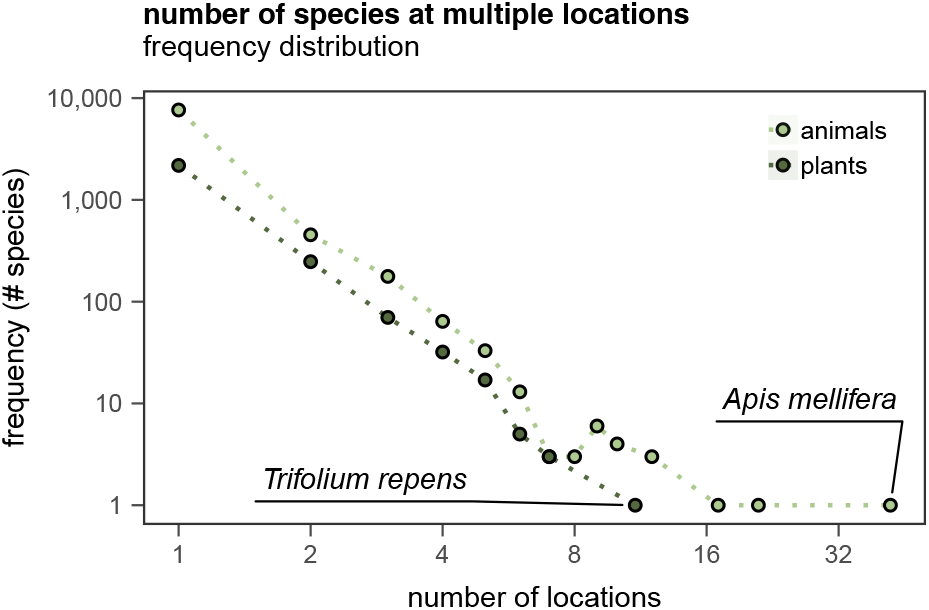
Frequency distribution of the number of locations in which a species is present. The most common pollinator species was *Apis mellifera*, which was sampled at 42 locations, while the most common plant species was *Trifolium repens*, which was sampled at 11 locations.

After matching species across studies as accurate as possible, we carried on two more steps. First, we calculated the environmental stress of species in their communities. Second, we relate the species stress in their community with the number of partner species it has as a metric of their level of specialisation.

### Environmental stress

We calculated the environmental stress of species in their communities. We assume that stress a species experiences in a particular location is inversely related to the suitability of the average environmental conditions in that place. As we aim to compare specialisation levels for different levels of environmental stress, we only calculate bioclimatic suitability for species that were present in at least two communities. To calculate the bioclimatic suitability of a species in a particular location, we used a niche-factor analysis (Hirzel et al. 2002; Broennimann et al. 2012). This approach is based on the probability density function of species distribution in an environmental variable space. Habitats are characterised by a collection of environmental variables. In a nutshell, those habitats in which the species occurs more often are deemed to be more suitable for the species than habitats in which the species has never been observed. As bioclimatic suitability is calculated in a scale from zero to one following the niche-factor analysis, for simplicity, we define environmental stress as one minus suitability.

The niche factor analysis requires two critical pieces of information. First, it requires information about the occurrences of the species of interest. Second, the method requires information about the environmental conditions for all the locations in which the species occurs. We retrieved 38.1 million occurrences from the Global Biodiversity Information Facility (GBIF; https://www.gbif.org). Issues with data quality are a central issue hampering the use of publicly available species occurrence GBIF data in ecology and biogeography (Jetz et al. 2019). We, therefore, followed a series of filters and geographic heuristics to correct or remove erroneous and imprecise referencing records (see supplementary methods; Zizka et al. 2019) which allowed us to identify and remove 7.5 million potentially problematic occurrences from further analysis. We integrated the occurrences from our plant-pollinator communities to the cleaned occurrences retrieved from GBIF.

We retrieved environmental data from WorldClim V2.0, which includes 19 bioclimatic variables commonly used in species distribution modelling (Fick and Hijmans 2017). We then complemented data obtained from WorldClim with data from Envirem (Title and Bemmels 2017), which includes 16 extra bioclimatic and two topographic variables. The additional set of variables from Envirem are relevant to ecological or physiological processes and thus have the potential to improve our suitability estimation (Title and Bemmels 2018). We obtained all environmental data as rasters composed by cells of 2.5 arc-minutes. We chose this resolution because it provides a reasonable match to the locational accuracy of the species occurrences found in GBIF, particularly those that originate from preserved specimens in museum collections.

After obtaining information about species occurrence and the environment, we then merged these two datasets such that a vector with details of our 37 bioclimatic and topographic variables characterised the location of each occurrence. Sets of occurrence data tend to be spatially aggregated due to sample bias (tendency to collect close to cities, certain countries). Moreover, spatial autocorrelation arises in ecological data because geographically clumped records tend to be more similar in physical characteristics and/or species abundances than do pairs of locations that are farther apart. To account for such spatial dependency in occurrence data, we only included one occurrence record if a species had more than one within a cell of the bioclimatic raster. We did this to avoid giving more weight to areas with a high number of occurrences, a common scenario in occurrence records collected opportunistically as the ones we use here. In this step we removed 85.4% of the occurrences which resulted in a total of 4.5 million occurrences used in our niche analysis.

A common issue of terrestrial bioclimatic datasets is that the boundaries of the cells with information do not precisely match the landmass boundaries. The result of this missmatch is that not all environmental variables were available for 3,273 of the raster cells with occurrences (0.8% of the total). As expected, the vast majority of these problematic cells were close to the shore. To address this issue, we calculated the average value of environmental variables within a 5km buffer of the centre of the cell where the variable was missing and used it to approximate the value of the variable in that cell. Using this procedure, we were able to fill environmental variables for 89.3% of the cells where they were missing. To fill the remaining 350 cells, we repeated the aforementioned procedure but instead using a 10km buffer. We removed from further analysis occurrences located within the 135 cells for which we were unable to fill environmental variables (0.03% of the total).

Next, we calculated the probability density function of the species distribution in environmental space. To determine the environmental space, we used the first two components from a principal component analysis of the 37 bioclimatic variables associated with the species occurrences. Specifically we used the `dudi.pca` function from the R package `ade4` 1.7.13 (Dray and Dufour 2007) and center and scale all bioclimatic variables to have a mean of zero and a unit variance. We then determined the position of species occurrences in the environmental space and estimate their bivariate probability density function. We used a kernel method to estimate this density and normalised it such that it ranges between zero and one. We used the kernel density method in the niche-factor analysis (Broennimann et al. 2012) rather than the distance from the mode (Hirzel et al. 2002) (as it has been proposed earlier) because it has been shown to reduce the procedure’s sensitivity to sampling effort and the resolution of the environmental space. Specifically, to calculate the probability density function we used `ecospat.grid.clim.dyn` from the R package `ecospat` 3.0 (Broennimann, Di Cola, and Guisan 2018) with a grid resolution of 200. We then determined the location in the environmental space of the plant-pollinator communities using the function `suprow` from `ade4`. The normalised density at that particular location (which we calculated using the R package `raster` 2.8.19; Hijmans 2019) corresponds the bioclimatic suitability. The result of all these steps is the environmental stress which corresponds to one minus the bioclimatic suitability for a species of a particular location.

We used a sensitivity analysis to determine the minimum number of occurrences that are necessary to have robust environmental stress estimations. For that we used the species with most occurrences available, *Archilochus colubris*, and calculated the mean absolute error of the bioclimatic suitability values obtained with one thousand subsamples from the 74,791 occurrences available from GBIF.

### Data analysis

We then used a set of Bayesian multilevel models to evaluate the impact of environmental stress on species specialisation. Specifically, we use the normalised degree of species as our response variable; that is, the number of species it interacts with given the number of species in the opposite guild (Martín González, Dalsgaard, and Olesen 2010). The normalised degree was modelled using a logit link function, and a binomial distribution in which the number of partner species a focal interacts with is the number of successes, and the number of species in the opposite guild is the number of trials. We are aware that whether species interact or not is not a Bernoulli process as species interactions are not strictly independent from each other. However, the use of a binomial distribution allows us to account for the differences in species richness across communities indirectly. Importantly, results are qualitatively similar when we model species degree directly using a Poisson distribution and a logarithmic link function.

We evaluated four models to assess the relative importance of suitability. A first model, our baseline model, included five variables. The predictors in the baseline model were the environmental stress, its number of known possible partners in the community, and both the species guild (plant or a pollinator) and its interaction with environmental stress. We included the number of known possible partners as a predictor in our models as it allows us to control for the effects of the environment on community composition, effectively accounting for species co-occurrence. We calculated this metric by determining the number of partners with which the species is known to interact in any other community. Controlling for the number of potential partners makes our model a particularly stringent test of our environmental-stress hypotheses because this variable could explain a large proportion of variance. Often, the potential and the actual number of partners is the same or very close to each other, especially for rare species present only in a few communities.

We allowed the intercept and slope of the stress-specialisation relationship to vary among species. This approach allowed us to investigate two questions. First, it allows us to inspect the extent to which environmental stress affects species similarly. Second, by investigating the correlation between the intercept and the slope as a model parameter, it allowed us to inspect the extent by which species with a small or large number of partner species respond to increasing levels of environmental stress. To account for unmeasured differences between communities, like sampling effort, sampling method, or diversity, we also allowed the model intercept to be different for each community in our study. To facilitate model interpretation and convergence, we scaled all continuous variables to have a mean of zero and a unit variance.

We compared this baseline model with three alternative models in which we removed one predictor at a time. To quantify the difference between models, in terms of their expected out-of-sample performance, we use the Wanatabe-Akaike information criterion (WAIC). All models were fitted under a Bayesian framework using the R package brms 2.8.0 (Bürkner 2017, 2018) as an interface for Stan (Carpenter et al. 2017). For each model, we used four Markov chains of 4,000 iterations each; we used half of the iterations for warmup. We used weakly informative priors for all model parameters. Specifically we used normal priors of mean zero and standard deviation ten for the population-level effects and the intercepts, a half-Cauchy prior with a location of zero and a scale of two for the standard deviations, and, when applicable, an LKJ-correlation prior with parameter *ζ* = 1 for the correlation matrix between group-level parameters.

## Results

After performing our sensitivity analysis, we found that, for a species, we need roughly 18 independent occurrences for each community for which we aim to estimate the environmental stress. This is the number of occurrences necessary to maintain the mean absolute error of bioclimatic suitability below 0.1 (Fig. 2). We therefore removed from further analyses 286 species for which we did not have enough occurrences to obtain robust estimates.

**Figure 2:**
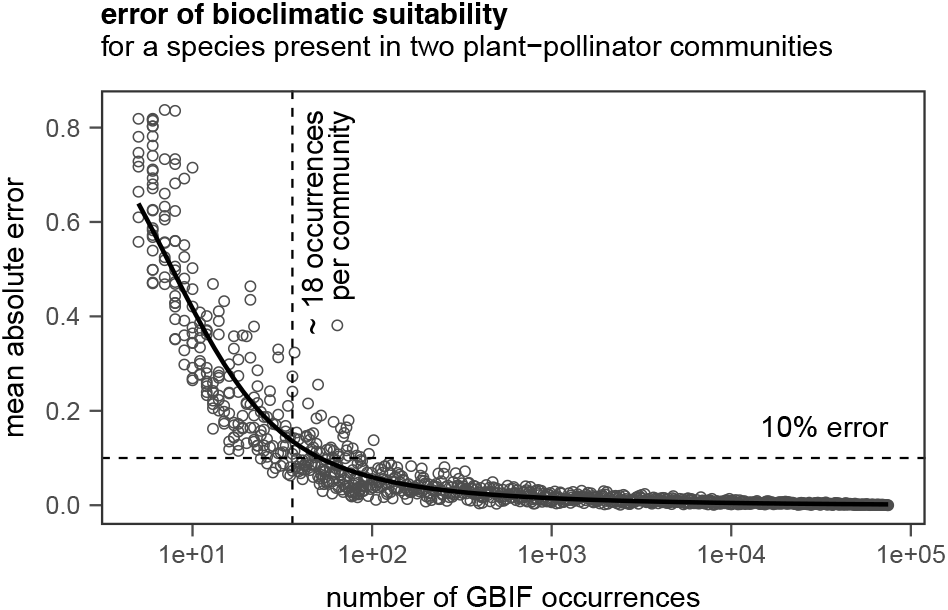
Sensitivity analysis of environmental stress error. The number of independent occurrences retrieved from GBIF is inversely related to the error of bioclimatic suitability for our plant-pollinator networks.The sensitivity analysis was performed by subsampling occurrences of *Archilochus colubris* the species in our dataset with the largest number of occurrences in GBIF, which was recorded in two of our communities.

Our models performed relatively well. The Bayesian R^2^ for our baseline model was 0.89, which indicates our models were able to capture a large proportion of the variability on the data. Overall, we found that environmental stress does not have a consistent effect across species. Indeed, when looking at the fixed effects, stress has virtually no relationship with the normalised degree—our metric of specialisation (Figure 3a). However, environmental stress was still an important predictor in our model. The difference in WAIC between our baseline model and the model that did not include environmental stress was 489 ± 94 (Table 1). This apparent discrepancy can be explained by the variability of the specialisation-stress relationship across species.

**Table 1:** Comparison in out of sample predictive power of the baseline model (bold) and their alternatives. We rank models by their expected log predictive density based on their Wanatabe-Akaike information criterion (WAIC).

| predictors | WAIC | SE |
| --- | --- | --- |
| **stress x guild + # possible partners** | 6,592 | 170 |
| stress + # possible partners | 6,595 | 166 |
| guild + # possible partners | 7,081 | 202 |
| stress x guild | 8,041 | 290 |

**Figure 3:**
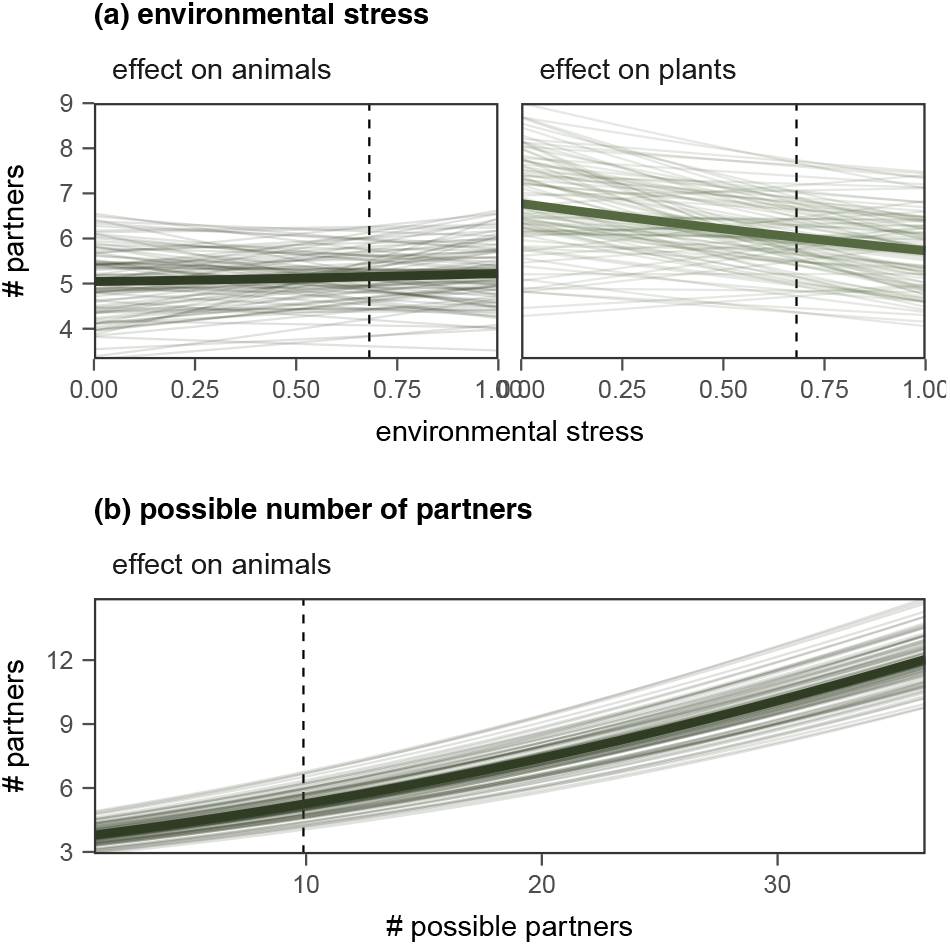
Conditional effects of predictors in our baseline model. The shown values are based on predictions for a hypothetical community with 76 plants and 33 pollinators. These values correspond to the median number of species in each guild across communities. In each panel, we condition on the mean value of the other predictor in the model. We indicate mean values for each predictor with a vertical dashed line. For model fitting, we scaled all predictors to have a mean of zero and unit variance; however, here we show the unscaled predictors to facilitate interpretation. To illustrate the uncertainty around the fitted estimates, we plot the fits of 100 independent draws from the posterior distribution. The thick lines indicate the mean values of the response distribution. As there was no interaction between the guild and the number of possible interactions, we only show the conditional effect of pollinators.

For some species, there is a strong negative relationship between stress and specialisation, while for others, there is a strong positive relationship (Figure 4a). Interestingly, the slope of this relationship correlates with the species’ intercept in the model (Figure 4b and c). Recall that the model estimates the intercept at the mean value for stress across communities (0.68). The mean correlation coefficient was 0.52 [0.33, 0.67]. Therefore, the slope of the stress-specialisation relationship was more likely to be positive for species with a large number of partners under average stress conditions (and negative for species with a smaller number of partners). Extrapolating to no-stress conditions: species that would interact with a small number of partners under no stress are more likely to interact with more partners as stress increases, whereas those that would interact with a large number of partners are more likely to interact with less.

**Figure 4:**
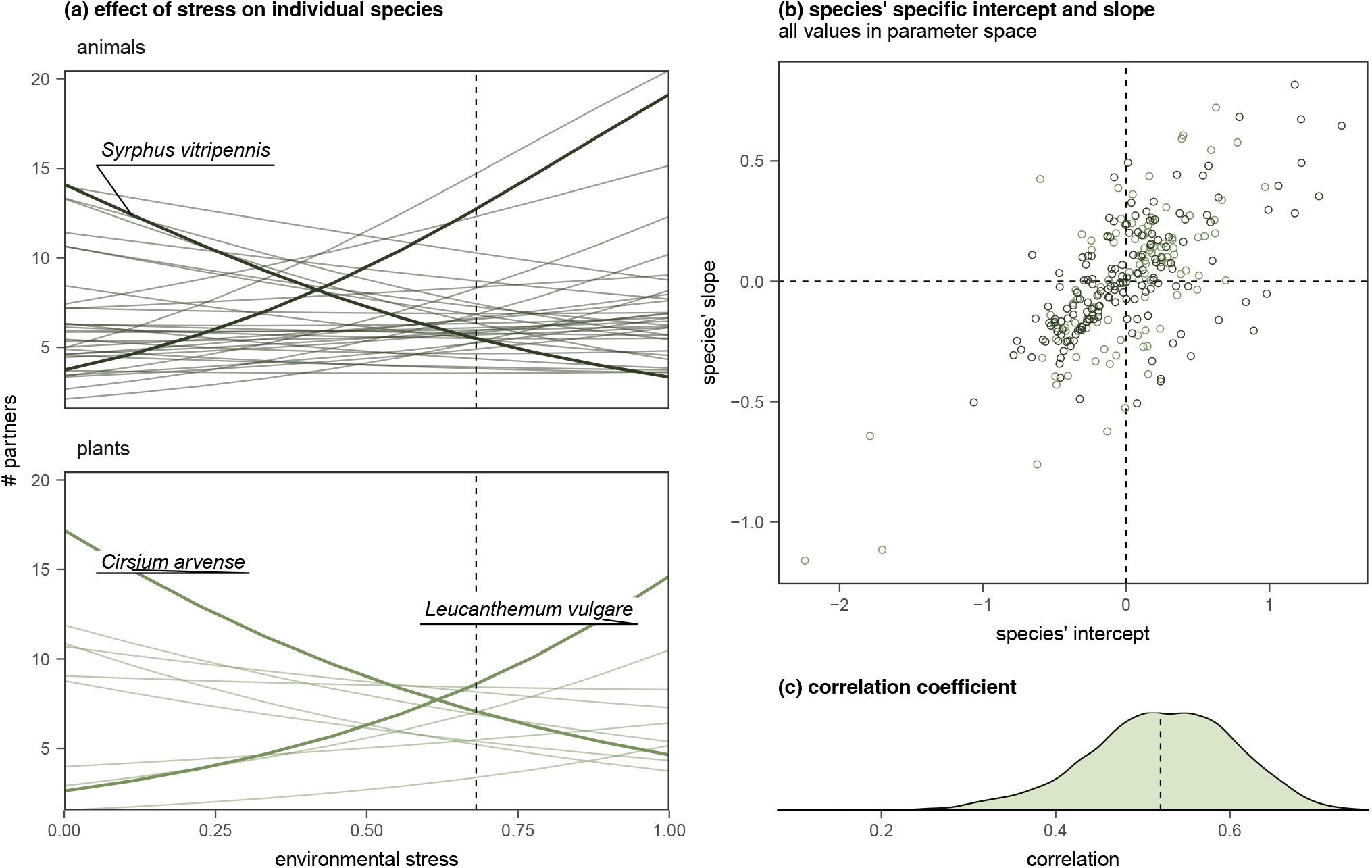
Species-level effects of environmental stress (a) Conditional effect of stress for individual species. Each line corresponds to the median relationship for each species. Although we included in the analysis of all species that are present in two or core communities, to facilitate visualisation here, we show only species for which there is suitability information in at least six communities (10 plants and 33 pollinators). As in the previous figure, fitted values assume a hypothetical community of median size. In each panel, we highlight two species for which the relationship between environmental suitability and the normalised degree was particularly strong. (b and c) The correlation between the species’ intercept and the species’ slope of suitability was negative. The species’ intercept can be interpreted as the relative difference between the number of partners a species has under mean levels of environmental stress and the mean number of partners across all species. Positive values of species’ slope indicate a positive relationship between stress and the number of partners and vice-versa.

As expected, we found a strong and positive relationship between the number of possible interactions and the number of realised interactions in the community. There was also a large difference of WAIC between the model that included this predictor and that that excluded it. This result indicates that the availability of potential partners—this is, community composition—accounts for a large proportion of the variability in species degree. Importantly, our findings relating to the variability of the stress-specialisation relationship were qualitatively unchanged, whether we included this variable or not.

The standard deviation (in the parameters scale) of the community intercepts was 1.02 [0.85, 1.23] which indicates the importance of the local context when determining specialisation. The standard deviation of the species intercept was 0.54 [0.48, 0.61], and that of the species’ stress slope was 0.38 [0.32, 0.44] (95% credible intervals shown within square brakets).

## Discussion

We set out to explore whether and how environmental stress can systematically affect specialisation. After accounting for the pool of potential partners, we found that environmental conditions contribute to determining whether a species is a generalist or a specialist *in their community*. We also found that the particular effect of the environment is strongly dependent on the species. Based on existing literature, we proposed two alternative hypotheses of how environmental stress may affect species’ specialisation, and we found evidence for both of them. Species with a large number of partners in low-stress communities were more likely to have a negative relationship and hence reduce the number of partners as stress increases. Contrastingly, species in our datasets with a small number of partners in low-stress communities were more likely to have a larger number of partners in more stressful communities. In summary, environmental stress pushes species that are flexible enough to change their interaction partners towards intermediate levels of specialisation, a so-called “regression towards the mean”.

Our results suggest that changes in community composition are indeed the primary channel through which the environment determines changes interaction probability. However, they also show that, for a large number of species, the environment may also play a substantial role in determining their level of specialisation. Previous research has recognised that environmental factors may help explain the changes in network structure along environmental gradients that cannot be explained by community composition (Tylianakis, Tscharntke, and Lewis 2007). However, how these two factors were linked had been elusive so far (Gravel et al. 2018). We believe that part of this difficulty could have arisen because species, and ultimately network structure, can respond in multiple, and contrasting, ways depending on the particular bioclimatic variable examined (e.g. temperature or precipitation). Using stress to summarise the effect on species of multiple environmental gradients allowed us to detect a clear signal of the environment in species’ interaction patterns.

Although both niche and neutral processes are relevant in determining species interactions, our model suggests that niche processes may be the predominant mechanism through which the environment *systematically* affects specialisation. First, it is unlikely that environmental stress correlates to local species abundances (Pearce and Ferrier 2001; Sagarin, Gaines, and Gaylord 2006). Second, even if there is a relationship between stress and abundances, a particular environmental gradient could have a positive effect on the abundance of some species and a negative effect on others. Indeed, we find that within a community there is a wide range of stress values, even for the relatively limited number of species we were able to include in our analysis.

Recent research suggests that species are continuously changing their interaction partners wherever environmental conditions change in space or time (Raimundo, Guimarães, and Evans 2018). So far it appears that this rewiring is primarily driven by generalist species (Ponisio, Gaiarsa, and Kremen 2017; Burkle, Marlin, and Knight 2013), presumably because generalist species are less sensitive to trait matching of their interaction partners (CaraDonna et al. 2017). Our results add two important nuances to these findings. First, because “generalists” seem to focus on a smaller number of partners as environmental conditions deteriorate, we show that trait matching might still play a role in determining the interactions of generalist species. Second, and most importantly, our results suggest that only a small proportion of species are “true generalists” or “true specialists” this is, species that interact with a large or small number of partners regardless of the environmental stress, respectively. This pattern implies that rewiring is not exclusive of species with a large number of partners. Instead, at least a fraction of the species that appear to be specialist *in their communities* might be as flexible, if not more, than those with a large number of partners, effectively behaving as facultative generalists in the face of environmental change. These flexible “specialists” might therefore have a more significant role in network persistence than previously expected.

In our model, we can roughly divide species between true specialists, true generalists, and flexible species. However, there is a fourth group that remained invisible to our model but has important implications for network persistence and stability. Species that can vary their interaction partners flexibly and their role in the network are more likely to persist in their community as environmental conditions vary (Gaiarsa, Kremen, and Ponisio 2019). We propose this fourth group is composed of true specialists that are constrained to interact with partners of high trait-matching and therefore were not likely to be found in more than one community. If species that are not flexible are unlikely to persist over temporal or spatial environmental gradients, we can expect specialised communities that are highly constrained by trait-matching (like some plant-hummingbird networks; Vizentin-Bugoni, Maruyama, and Sazima 2014; Maruyama et al. 2014) to be far more vulnerable to increased climate change-induced environmental stress and habitat degradation than communities where role and interaction flexibility are more prevalent.

Similarly, if the patterns we see in our models have also played a role during the evolutionary history of pollination communities, our results also help explain why only a small fraction of plant-pollinator interactions show a strong signature of deep co-evolutionary history (Hutchinson, Cagua, and Stouffer 2017). The increases in the stress that species are predicted to experience due to rapid environmental change might further erode the co-evolutionary history of specialist species. Communities as a whole might be in a trajectory of even more diffuse co-evolution. For specialists, at least, the longer-term benefits of being able to interact with multiple partners might be more important than the shorter-term benefits of interacting with partners of high trait matching.

The structural implications of the “regression towards the mean” that environmental stress promotes are less clear. However, it is plausible to expect that nestedness, and therefore network stability, might be reduced in the face of rapid environmental change. Determining exactly how the changes in degree caused by environmental stress reflect on systematic changes in network structure would be an interesting avenue of research. Answering this question would require expanding our suitability analysis to all species in the community and compare the degree distribution of networks along a gradient of stress for the community as a whole.

In conclusion, we show that the environment can affect the specialisation level of plants and pollinators in systematic ways beyond community composition. Species that are inflexible with their interaction partners are unlikely to persist under more stressful environmental conditions. However, we show that many species are flexible in regards to their specialisation levels and therefore are not inherently generalists or specialists. Instead, the species’ level of specialisation/generalisation should be considered on a relative scale depending on where they are found and the environmental conditions at that location.

## Acknowledgements

We thank Warwick Allen, Marilia Gaiarsa, and Guadalupe Peralta for feedback and valuable discussions. EFC acknowledges the support from the University of Canterbury Doctoral Scholarship and a New Zealand International Doctoral Research Scholarship administered by New Zealand Education. DBS and JMT acknowledge the support of Rutherford Discovery Fellowships (RDF-13-UOC-003 and RDF-UOC-1002) and the Marsden Fund Council (UOC-1705), administered by the Royal Society of New Zealand Te Apārangi.

