## Supplementary Information for "Environment affects specialisation of plants and pollinators"

*E. Fernando Cagua, Audrey Lustig, Jason M. Tylianakis, Daniel B. Stouffer*

### Supplementary methods

#### Reducing taxonomic uncertainty

Data were obtained from the Web of Life database (Fortuna, Ortega, and Bascompte 2014) which includes data from 57 published studies (Abreu and Vieira 2004; Arroyo, Primack, and Armesto 1982; Barrett and Helenurm 1987; Bartomeus, Vilà, and Santamaría 2008; Bek 2006; Bezerra, Machado, and Mello 2009; Bundgaard 2003; Canela 2006; Clements and Long 1923; del Coro Arizmendi and Ornelas 1990; Dicks, Corbet, and Pywell 2002; Dupont and Olesen 2009; Dupont, Hansen, and Olesen 2003; Elberling and Olesen 1999; Gutierrez, Rojas-Nossa, and Stiles 2004; Hattersley-Smith 1985; Herrera 1988; Hocking 1968; Ingversen 2006; INouE et al. 1990; Inouye and Pyke 1988; Kaiser-Bunbury et al. 2014, 2010; Kakutani et al. 1990; Kato 2000; Kato, Matsumoto, and Kato 1993; Kato and Miura 1996; Kato et al. 1990; Kevan 1970; Kohler 2011; Lara 2006; Las-Casas, Azevedo Júnior, and Dias Filho 2012; Lundgren and Olesen 2005; McMullen 1993; Medan et al. 2002; Memmott 1999; Montero 2005; Mosquin 1967; Motten 1986; Olesen, Eskildsen, and Venkatasamy 2002; Ollerton 2003; Percival 1974; Petanidou and Vokou 1993; Philipp et al. 2006; Primack 1983; Ramirez 1989; Ramirez and Brito 1992; Robertson 1929; Rosero and others 2003; Sabatino 2010; Schemske et al. 1978; Small 1976; Smith-Ramírez et al. 2005; Stald, Valido, and Olesen 2003; Vázquez 2002; Vizentin-Bugoni et al. 2016; Yamazaki and Kato 2003).

Interaction data from the included studies included 11,231 unique organism names. From these 1,166 were present in more than one study. From the total number of organisms, 159 were identified to the subspecies or variety level, 6,759 to the species level, 1,755 to the genus level, whereas the remaining 2,558 were unidentified. As the species level was the most common taxonomic rank available in our interaction datasets, in all further analysis, we grouped together subspecies or varieties within the same species.

We were able to confirm the validity of 5,263 of the scientific names used to identify organisms (roughly 76%). We assessed the validity of a name by querying the Global Names Resolver database (https://resolver.globalnames.org) which includes data from 98 taxonomic sources. We accessed this database using the function `gnr_resolve` from the R package `taxize` 0.9.6 (Chamberlain and Szocs 2013; S. Chamberlain, Szoecs, et al. 2019).

From the remaining 1,655 names we were unable to validate, we were able to identify and correct 726 that contained spelling mistakes. These spelling mistakes were corrected automatically by fuzzy matching the canonical names in our data sources with those in the Global Names Resolver database. However, on rare occasions, the fuzzy matching algorithm can suggest a scientific name that has a similar spelling, but that corresponds to an organism in a different taxonomic group, often a separate kingdom. To address this potential problem, we checked the taxonomic hierarchy of suggested names and confirmed that it matched our expected taxon. We retrieved all taxonomic hierarchies from the National Center for Biotechnology Information taxonomic database (https://www.ncbi.nlm.nih.gov/taxonomy).

As species names are constantly changing, we subsequently checked for possible synonyms of the canonical names in our data sources. Using data from the Integrated Taxonomic Information System database (http://www.itis.gov), we found synonyms and alternative names for 611 species.
Finding these alternative names was required for two main reasons. First, because we wanted to be able to identify the cases in which the same species might have been recorded with different names in various data sources. This can occur not only when the canonical name has been changed but also when there are widely used orthographic variants. Second, because retrieving occurrence data is often only possible using the latest accepted/valid name for a particular species.

All together, from the 1,655 names we were unable to validate, it was not possible to automatically correct or find synonyms 332 of them. We then manually consulted multiple online databases, chiefly Wikispecies (https://species.wikimedia.org/), and looked for canonical names that both, resembled the unvalidated names and matched the geographic and taxonomic expectations. In this fashion, we were able to further correct 25 names. Most manual corrections were made on names that have been abbreviated or had more than two spelling mistakes. A complete list of manual name corrections can bee seen in Table S1.

This cleaning process allowed us to match further 270 names across data

Table S1: Manually corrected canonnical names. More than one correct name have been included when an accepted/valid synonym the cannonical name exists.

| incorrect name | corrected name | guild |
|---|---|---|
| Acaena pinn | *Acaena pinnatifida* | plant |
| Adesmia brachy | *Adesmia brachysemeon* | plant |
| Aesculus camea | *Aesculus X carnea* | plant |
| Brachyome sinclairii | *Brachyscome sinclairii* | plant |
| Calceolaria arac | *Calceolaria arachnoidea* | plant |
| Equium sabulicola | *Echium sabulicola* | plant |
| Euonymus fo rtunei | *Euonymus fortunei* | plant |
| Galvezia leucantha pubescen | *Galvezia leucantha* | plant |
| Heliconia simulans | *Heliconia angusta* | plant |
| Pitcaimia flammea | *Pitcairnia flammea* | plant |
| Psittacanthus flavo viridis | *Psittacanthus flavo-viridis* | plant |
| Rodophiala bifidum | *Rhodophiala bifida* | plant |
| Stachys albi | *Stachys albicaulis* | plant |
| Stenactis annuus | *Erigeron annuus* | plant |
| Thaspium aureum atropurpurem | *Thaspium trifoliatum* | plant |
| Tristhema mauritiana | *Tristemma mauritianum* | plant |
| Tropaeolum polyph | *Tropaeolum polyphyllum* | plant |
| Tyttnera scabra | *Turnera scabra* | plant |
| | *Turnera ulmifolia* | plant |
| VVedelia biflora | *Melanthera biflora* | plant |
| | *Wedelia biflora* | plant |
| Cateres pennatus | *Kateretes pennatus* | pollinator |
| Eclimus harrisi | *Condylostylus crinicauda* | pollinator |
| Ptilandrena g. maculati | *Andrena distans* | pollinator |
| Tapinotaspis caerulea | *Chalepogenus caeruleus* | pollinator |
| Tapinotaspis herbsti | *Chalepogenus herbsti* | pollinator |

sources and, by doing so, identify another 72 species that were present in more than one study. The process also allowed us to identify problematic data sources in which some names were included as both plants and pollinators. These data sources were removed from further analysis. In seven of our data sources interaction data was recorded at multiple points in time. When this was the case, we combined interaction data into one single interaction network.

### Reducing location uncertainty

We retrieved occurrences from the Global Biodiversity Information Facility (GBIF; https://www.gbif.org) using the R package `rgbif` 0.9.6 (Chamberlain and Boettiger 2017; S. Chamberlain, Barve, et al. 2019). Specifically, for each species, we only requested occurrences for which the coordinates of the observation were available and that had no known geospatial issue in the GBIF database. Roughly, we downloaded 38.1 million occurrences for the 986 species we were interested on. This occurrences, however, contain observations of mixed quality. Therefore, we followed Zizka et al. (2019) and

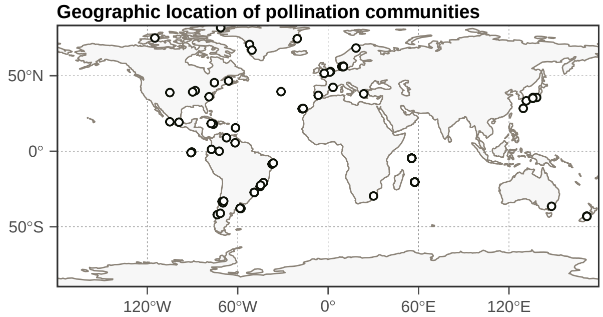


Figure S1: Worldwide distribution of pollination communities included in this study

applied a series of simple filters and geographic heuristics to remove those of lower quality. Specifically, we removed all occurrences with (*i*) a coordinate uncertainty larger than 100km; (*ii*) those recorded prior to 1945 (as records prior to this date have been shown to be often imprecise); (*iii*) those in which the number of counts in the occurrence was registered was either zero (as that indicates that the species has not been recorded); and (*iv*) those occurrences in which the "basis of record" was not a human observation or a preserved specimen (as occurrences from unknown and fossil records are known to be highly unreliable). We then used the R package `CoordinateCleaner` 0.9.6 (Zizka et al. 2019) and land mass and country data from Natural Earth (https://www.naturalearthdata.com) with a 1:10,000,000 scale to further identify and remove problematic occurrences. We removed occurrences for which their coordinates (*v*) fell outside the borders of the country where they were recorded; (*vi*) those around a country capital or the centroid of the country and province centroids; (*vii*) those around a biodiversity institution; and (*viii*) those located within oceans. Through this cleaning process, we removed with 7.5 million occurrences distributed across 916 species.

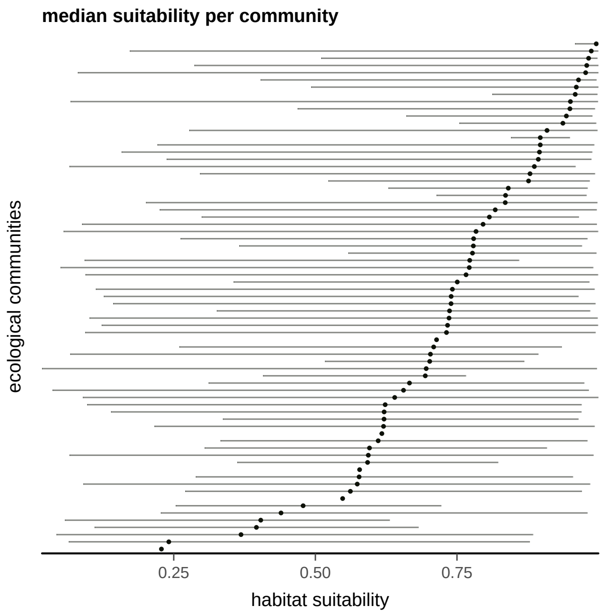


Figure S2: Median habitat suitability of communities in our dataset. Each row represents a different community and horizontal lines represent span the 2.5 and 97.5 quantiles.

Barrett, Spencer C. H., and Kaius Helenurm. 1987. "The Reproductive Biology of Boreal Forest Herbs. I. Breeding Systems and Pollination." *Canadian Journal of Botany* 65 (10): 2036–46. https://doi.org/10.1139/b87-278.

Bartomeus, Ignasi, Montserrat Vilà, and Luís Santamaría. 2008. "Contrasting Effects of Invasive Plants in Plant-Pollinator Networks." *Oecologia* 155 (4): 761–70. https://doi.org/10.1007/s00442-007-0946-1.

Bek, S. 2006. "A Pollination Network from a Danish Forest Meadow." MSc Thesis, Aarhus, Denmark: Aarhus University.

Bezerra, Elisângela L.S., Isabel C. Machado, and Marco A. R. Mello. 2009. "Pollination Networks of Oil-Flowers: A Tiny World Within the Smallest of All Worlds: Pollination Networks of Oil-Flowers." *Journal of Animal Ecology* 78 (5): 1096–1101. https://doi.org/10.1111/j.1365-2656.2009.01567.x.

Bundgaard, M. 2003. "Tidslig Og Rumlig Variation I et Plante-Bestøvernetv\a Erk." MSc Thesis, Aarhus, Denmark: Aarhus University.

Canela, Maria Bernadete. 2006. "Interações Entre Plantas E Beija-Flores Numa Comunidade de Floresta Atlantica Montana Em Itatiaia, Rio de Janeiro." PhD Thesis, Campinas, Brazil: Universidade Estadual de Campinas.

Chamberlain, Scott A, and Carl Boettiger. 2017. "R Python, and Ruby

Clients for GBIF Species Occurrence Data." Preprint. PeerJ Preprints. https://doi.org/10.7287/peerj.preprints.3304v1.

Chamberlain, Scott, Vijay Barve, Dan Mcglinn, Damiano Oldoni, Peter Desmet, Laurens Geffert, and Karthik Ram. 2019. *Rgbif: Interface to the Global Biodiversity Information Facility API.*

Chamberlain, Scott, and Eduard Szocs. 2013. "Taxize - Taxonomic Search and Retrieval in R." *F1000Research.*

Chamberlain, Scott, Eduard Szoecs, Zachary Foster, Zebulun Arendsee, Carl Boettiger, Karthik Ram, Ignasi Bartomeus, et al. 2019. *Taxize: Taxonomic Information from Around the Web.*

Clements, Frederic Edward, and Frances Louise Long. 1923. *Experimental Pollination: An Outline of the Ecology of Flowers and Insects.* 336. Carnegie Institution of Washington.

del Coro Arizmendi, Ma., and Juan Francisco Ornelas. 1990. "Hummingbirds and Their Floral Resources in a Tropical Dry Forest in Mexico." *Biotropica* 22 (2): 172. https://doi.org/10.2307/2388410.

Dicks, L. V., S. A. Corbet, and R. F. Pywell. 2002. "Compartmentalization in Plant-Insect Flower Visitor Webs." *Journal of Animal Ecology* 71 (1): 32–43. https://doi.org/10.1046/j.0021-8790.2001.00572.x.

Dupont, Yoko L., Dennis M. Hansen, and Jens M. Olesen. 2003. "Structure of a Plant-Flower-Visitor Network in the High-Altitude Sub-Alpine Desert of Tenerife, Canary Islands." *Ecography* 26 (3): 301–10. https://doi.org/10.1034/j.1600-0587.2003.03443.x.

Dupont, Yoko L., and Jens M. Olesen. 2009. "Ecological Modules and Roles of Species in Heathland Plant-Insect Flower Visitor Networks." *Journal of Animal Ecology* 78 (2): 346–53. https://doi.org/10.1111/j.1365-2656.2008.01501.x.

Elberling, Heidi, and Jens M. Olesen. 1999. "The Structure of a High Latitude Plant-Flower Visitor System: The Dominance of Flies." *Ecography* 22 (3): 314–23. https://doi.org/10.1111/j.1600-0587.1999.tb00507.x.

Fortuna, Miguel A., Raul Ortega, and Jordi Bascompte. 2014. "The Web of Life." *arXiv:1403.2575 [Q-Bio]*, March. http://arxiv.org/abs/1403.2575.

Gutierrez, Aquiles, Sandra Victoria Rojas-Nossa, and Gary F Stiles. 2004. "DINÁMICA ANUAL DE LA INTERACCIÓN COLIBRÍ-FLOR EN ECOSISTEMAS ALTOANDINOS." *Ornitologia Neotropical* 15: 205–13.

Hattersley-Smith, G. 1985. “Botanical Studies in the Lake Hazen Region, Northern Ellesmere Island, Northwest Territories, Canada, by James H. Soper and John M. Powell.” *ARCTIC* 38 (4): 346. https://doi.org/10.14430/arctic2421.

Herrera, Javier. 1988. “Pollination Relationships in Southern Spanish Mediterranean Shrublands.” *The Journal of Ecology* 76 (1): 274. https://doi.org/10.2307/2260469.

Hocking, Brian. 1968. “Insect-Flower Associations in the High Arctic with Special Reference to Nectar.” *Oikos* 19 (2): 359. https://doi.org/10.2307/3565022.

Ingversen, TT. 2006. “Plant-Pollinator Interactions on Jamaica and Dominica: The Centrality, Asymmetry and Modularity of Networks.” MSc Thesis, Aarhus, Denmark: University of Aarhus.

INouE, Tamiji, Makoto KATo, Takehiko KAKuTANi, Takeshi SuKA, and Takao It. 1990. “Of Kibune, Kyoto: An Overview of the Flowering Phenology and the Seasonal Pattern of Insect Visits’,” 88.

Inouye, David W., and Graham H. Pyke. 1988. “Pollination Biology in the Snowy Mountains of Australia: Comparisons with Montane Colorado, USA.” *Austral Ecology* 13 (2): 191–205. https://doi.org/10.1111/j.1442-9993.1988.tb00968.x.

Kaiser-Bunbury, Christopher N., Stefanie Muff, Jane Memmott, Christine B. Müller, and Amedeo Caflisch. 2010. “The Robustness of Pollination Networks to the Loss of Species and Interactions: A Quantitative Approach Incorporating Pollinator Behaviour.” *Ecology Letters* 13 (4): 442–52. https://doi.org/10.1111/j.1461-0248.2009.01437.x.

Kaiser-Bunbury, Christopher N., Diego P. Vázquez, Martina Stang, and Jaboury Ghazoul. 2014. “Determinants of the Microstructure of PlantPollinator Networks.” *Ecology* 95 (12): 3314–24. https://doi.org/10.1890/14-0024.1.

Kakutani, Takehiko, Tamiji Inoue, Makoto Kato, and Hidekuji Ichihashi. 1990. “Insect-Flower Relationship in the Campus of Kyoto University, Kyoto : An Overview of the Flowering Phenology and the Seasonal Pattern of Insect Visits.” *Contributions from the Biological Laboratory, Kyoto University* 27 (4): 465–522.

Kato, Makoto. 2000. “Anthophilous Insect Community and Plant-Pollinator Interactions on Amami Islands in the Ryukyu Archipelago, Japan,” 101.

Kato, Makoto, Takehiko Kakutani, Tamiji Inoue, and Takao Itino. 1990. "Insect-Flower Relationship in the Primary Beech Forest of Ashu, Kyoto: An Overview of the Flowering Phenology and the Seasonal Pattern of Insect Visits," 68.

Kato, Makoto, Masamichi Matsumoto, and Toru Kato. 1993. "Flowering Phenology and Anthophilous Insect Community in the Cool-Temperate Subalpine Forests and Meadows at Mt. Kushigata in the Central Part of Japan," 58.

Kato, Makoto, and Reiichi Miura. 1996. "Flowering Phenology and Anthophilous Insect Community at a Threatened Natural Lowland Marsh at Nakaikemi in Tsuruga, Japan." *Contributions from the Biological Laboratory, Kyoto University* 29 (1): 1.

Kevan, PG. 1970. "High Arctic Insect-Flower Relations: The Interrelationships of Arthropods and Flowers at Lake Hazen, Ellesmere Island, Northwest Territories, Canada." PhD Thesis, Edmonton, Canada: University of Alberta.

Kohler, Glauco Ubiratan. 2011. "Redes de Interação Planta Beija-Flor Em Um Gradiente Altitudinal de Floresta Atlântica No Sul Do Brasil." MSc Thesis, Curitiba, Brasil: Universidade Federal do Paraná.

Lara, Carlos. 2006. "Temporal Dynamics of Flower Use by Hummingbirds in a Highland Temperate Forest in Mexico." *Ecoscience* 13 (1): 23–29. https://doi.org/10.2980/1195-6860(2006)13[23:TDOFUB]2.0.CO;2.

Las-Casas, Fmg, Sm Azevedo Júnior, and Mm Dias Filho. 2012. "The Community of Hummingbirds (Aves: Trochilidae) and the Assemblage of Flowers in a Caatinga Vegetation." *Brazilian Journal of Biology* 72 (1): 51–58. https://doi.org/10.1590/S1519-69842012000100006.

Lundgren, Rebekka, and Jens M. Olesen. 2005. "The Dense and Highly Connected World of Greenland's Plants and Their Pollinators." *Arctic, Antarctic, and Alpine Research* 37 (4): 514–20. https://doi.org/10.1657/1523-0430(2005)037[0514:TDAHCW]2.0.CO;2.

McMullen, C. K. 1993. "Flower-Visiting Insects of the Galapagos Islands." *The Pan-Pacific Entomologist* 69 (1): 95.

Medan, Diego, Norberto H. Montaldo, Mariano Devoto, Anita Mantese, Viviana Vasellati, German G. Roitman, and Norberto H. Bartoloni. 2002. "Plant-Pollinator Relationships at Two Altitudes in the Andes of Mendoza, Argentina." *Arctic, Antarctic, and Alpine Research* 34 (3): 233. https://doi.org/10.2307/1552480.

Memmott, J. 1999. "The Structure of a Plant-Pollinator Food Web." *Ecology Letters* 2 (5): 276–80. https://doi.org/10.1046/j.1461-0248.1999.00087.x.

Montero, AC. 2005. "The Ecology of Three Pollination Networks." MSc Thesis, Aarhus, Denmark: Aarhus University.

Mosquin, Theodore. 1967. "Observations on the Pollination Biology of Plants on Melville Island, NWT." *Can. Fld Nat.* 81: 201–5.

Motten, Alexander F. 1986. "Pollination Ecology of the Spring Wildflower Community of a Temperate Deciduous Forest." *Ecological Monographs* 56 (1): 21–42. https://doi.org/10.2307/2937269.

Olesen, Jens M., Louise I. Eskildsen, and Shadila Venkatasamy. 2002. "Invasion of Pollination Networks on Oceanic Islands: Importance of Invader Complexes and Endemic Super Generalists." *Diversity <Html_ent Glyph="@amp;" Ascii="&"/> Distributions* 8 (3): 181–92. https://doi.org/10.1046/j.1472-4642.2002.00148.x.

Ollerton, J. 2003. "The Pollination Ecology of an Assemblage of Grassland Asclepiads in South Africa." *Annals of Botany* 92 (6): 807–34. https://doi.org/10.1093/aob/mcg206.

Percival, Mary. 1974. "Floral Ecology of Coastal Scrub in Southeast Jamaica." *Biotropica* 6 (2): 104. https://doi.org/10.2307/2989824.

Petanidou, Theodora, and Despina Vokou. 1993. "POLLINATION ECOLOGY OF LABIATAE IN A PHRYGANIC (EAST MEDITERRANEAN) ECOSYSTEM." *American Journal of Botany* 80 (8): 892–99. https://doi.org/10.1002/j.1537-2197.1993.tb15310.x.

Philipp, Marianne, Jens Böcher, Hans R. Siegismund, and Lene R. Nielsen. 2006. "Structure of a Plant-Pollinator Network on a Pahoehoe Lava Desert of the Galápagos Islands." *Ecography* 29 (4): 531–40. https://doi.org/10.1111/j.0906-7590.2006.04546.x.

Primack, Richard B. 1983. "Insect Pollination in the New Zealand Mountain Flora." *New Zealand Journal of Botany* 21 (3): 317–33. https://doi.org/10.1080/0028825X.1983.10428561.

Ramirez, Nelson. 1989. "Biologia de Polinizacion En Una Comunidad Arbustiva Tropical de La Alta Guayana Venezolana." *Biotropica* 21 (4): 319. https://doi.org/10.2307/2388282.

Ramirez, Nelson, and Ysaleny Brito. 1992. "Pollination Biology in a Palm Swamp Community in the Venezuelan Central Plains." *Botanical Journal of* *the Linnean Society* 110 (4): 277–302. https://doi.org/10.1111/j.1095-8339.1992.tb00294.x.

Robertson, Charles. 1929. *Flowers and Insects Lists of Visitors of Four Hundred and Fifty Three Flowers.* Cairnville: The Science Press Printing Company.

Rosero, Liliana, and others. 2003. "Interações Planta/Beija-Flor Em Três Comunidades Vegetais Da Parte Sul Do Parque Nacional Natural Chiribiquete, Amazonas (Colômbia)." PhD Thesis, Campinas, Brazil: Universidade Estadual de Campinas.

Sabatino, Malena. 2010. "Direct Effects of Habitat Area on Interaction Diversity in Pollination Webs." *Ecological Applications* 20 (6): 7.

Schemske, Douglas W., Mary F. Willson, Michael N. Melampy, Linda J. Miller, Louis Verner, Kathleen M. Schemske, and Louis B. Best. 1978. "Flowering Ecology of Some Spring Woodland Herbs." *Ecology* 59 (2): 351–66. https://doi.org/10.2307/1936379.

Small, Ernest. 1976. "Insect Pollinators of the Mer Bleue Peat Bog of Ottawa." *Canadian Field-Naturalist.*

Smith-Ramírez, C., P. Martinez, M. Nuñez, C. González, and J. J. Armesto. 2005. "Diversity, Flower Visitation Frequency and Generalism of Pollinators in Temperate Rain Forests of Chiloé Island, Chile." *Botanical Journal of the Linnean Society* 147 (4): 399–416. https://doi.org/10.1111/j.1095-8339.2005.00388.x.

Stald, L, A Valido, and Jens M. Olesen. 2003. "Struktur Og Dynamik I Rum Og Tid at et Bestøvningsnetværk På a Tenerife, De Kanariske Øer." MSc-Thesis, Aarhus, Denmark: University of Aarhus.

Vázquez, Diego P. 2002. "Interactions Among Introduced Ungulates, Plants, and Pollinators: A Field Study in the Temperate Forest of the Southern Andes." PhD Thesis, Knoxville, United States of America: University of Tennessee.

Vizentin-Bugoni, Jeferson, Pietro K. Maruyama, Vanderlei J. Debastiani, L. da S. Duarte, Bo Dalsgaard, and Marlies Sazima. 2016. "Influences of Sampling Effort on Detected Patterns and Structuring Processes of a Neotropical Plant-Hummingbird Network." Edited by Daniel B. Stouffer. *Journal of Animal Ecology* 85 (1): 262–72. https://doi.org/10.1111/1365-2656.12459.

Yamazaki, Kyoko, and Makoto Kato. 2003. "Flowering Phenology and

Anthophilous Insect Community," 67.

Zizka, Alexander, Daniele Silvestro, Tobias Andermann, Josué Azevedo, Camila Duarte Ritter, Daniel Edler, Harith Farooq, et al. 2019. "CoordinateCleaner: Standardized Cleaning of Occurrence Records from Biological Collection Databases." Edited by Tiago Quental. *Methods in Ecology and Evolution* 10 (5): 744–51. https://doi.org/10.1111/2041-210X.13152.
